# HIV-1-induced apathy: Mitigation by the gut metabolite, S-Equol

**DOI:** 10.1101/2021.01.04.425260

**Authors:** Kristen A. McLaurin, Sarah J. Bertrand, Jessica M. Illenberger, Steven B. Harrod, Charles F. Mactutus, Rosemarie M. Booze

**Affiliations:** Program in Behavioral Neuroscience, Department of Psychology, Barnwell College, 1512 Pendleton Street, University of South Carolina, Columbia, SC 29208

**Keywords:** Apathy, HIV-1 Associated Neurocognitive Disorders, S-Equol, Cocaine Self-Administration, Estrogen

## Abstract

The persistence of motivational alterations, including apathy, in older HIV-1 seropositive individuals, despite treatment with combination antiretroviral therapy, necessitates the development of innovative adjunctive therapeutics. S-Equol (SE), a selective estrogen receptor β agonist, has been implicated as a neuroprotective and/or neurorestorative therapeutic for HIV-1 associated neurocognitive disorders (HAND); its therapeutic utility for apathy, however, has yet to be systematically evaluated. Thus, beginning at approximately seven to nine months of age, HIV-1 transgenic (Tg) and control animals were treated with either a daily oral dose of SE (0.2 mg) or vehicle and assessed in a series of tasks to evaluate goal-directed behavior. First, at the genotypic level, apathetic behavior in older HIV-1 Tg rats treated with vehicle was characterized by a diminished reinforcing efficacy of, and sensitivity to, sucrose and enhanced drug seeking for cocaine relative to control animals treated with vehicle. Second, treatment with SE ameliorated alterations in goal-directed behaviors and reduced drug seeking behavior in HIV-1 Tg rats. Following a history of cocaine self-administration, HIV-1 Tg animals treated with vehicle exhibited prominent decreases in dendritic branching and a shift towards longer dendritic spines with decreased head diameter; synaptic dysfunction that was partially restored by SE treatment. Taken together, SE restored motivated behavior in the HIV-1 Tg rat, expanding the potential clinical utility of SE to include both neurocognitive and affective alterations.

## INTRODUCTION

The great success of combination antiretroviral therapy (cART) has led to a growing population of older individuals (>50 years of age) living with human immunodeficiency virus type 1 (HIV-1). Indeed, worldwide, approximately 7.9 million HIV-1 seropositive individuals are aged 50 and older, accounting for approximately 20.7% of HIV-1 seropositive individuals [1]; a prevalence that is significantly higher in high-resource countries (i.e., approximately 51% in the United States, [2]). However, despite achieving near-normal life expectancy [3], older HIV-1 seropositive individuals exhibit an increased neurobehavioral burden, including more frequent apathy, relative to their seronegative counterparts [4].

Apathy, which has been recognized as a key neuropsychiatric feature of HIV-1 since early in the epidemic [5], afflicts between 26-42% of HIV-1 seropositive individuals in the post-cART era [6-7]. Recognized as a multidimensional syndrome resulting in diminished motivation [8], apathy is characterized by a quantitative reduction in self-generated voluntary and purposeful (goal-directed) behaviors [9]. The severity of apathy is associated with profound functional consequences, including greater impairment in activities of daily living [7, 10-11], an increased number of cognitive complaints [7,10], decreased medication adherence [12-13], and decreased health-related quality of life [14]. Furthermore, amongst HIV-1 seropositive individuals, apathy is positively correlated with both disease duration [15] and age [16]. Thus, there remains a critical need to address apathy following chronic HIV-1 viral protein exposure and develop innovative adjunctive therapeutics to target the prominent neurobehavioral deficits in aging HIV-1 seropositive individuals.

The anterior cingulate circuit, one of the five frontal-subcortical circuits linking the basal ganglia and prefrontal cortex (PFC; [e.g., 17]), plays a critical role in motivational mechanisms and goal-directed behavior [18]. Broadly, the anterior cingulate circuit is comprised of the anterior cingulate cortex, the ventral striatum, including the nucleus accumbens (NAc), and basal ganglia [17]. More specifically, medium spiny neurons (MSNs) of the NAc play a functional role in many motivational [e.g., 19-20] and goal-directed [e.g., 21] behaviors. MSNs are characterized by a centrifugal morphology and high densities of dendritic spines [22], which receive excitatory inputs [23] and may reflect neuronal processing capacity [24]. Chronic HIV-1 viral protein exposure damages MSNs evidenced by alterations in dendritic branching complexity [25-26], prominent synaptic dysfunction [25-27], and dysregulation of neuronal excitability [28]. Furthermore, MSNs have been implicated as a key structural loci for the actions of HIV-1 viral proteins on goal-directed behaviors [26]; a loci which affords a key target for the development of innovative therapeutics.

Equol, a phytoestrogen that is structurally similar to 17β-estradiol [29], may serve as a novel therapeutic to target the prominent synaptic dysfunction observed in MSNs following chronic HIV-1 viral protein exposure. Following the ingestion of the soy derived phytoestrogen daidzein, Equol is produced by the gut microbiota [30]. S-Equol (SE), the only enantiomer produced by humans [31], penetrates the central nervous system via the blood-brain barrier and exhibits selective affinity for estrogen receptor β (ERβ; [31-32]). With regards to HIV-1, pretreatment with SE precludes synapse loss resulting from exposure to the HIV-1 viral protein, Tat [32]; research which corroborates studies demonstrating the utility of daidzein, a precursor to SE [30], to both protect and restore synaptodendritic damage in HIV-1 [33]. Furthermore, SE has been implicated as an efficacious therapeutic for the neurocognitive impairments, collectively termed HIV-1 associated neurocognitive disorders (HAND), associated with the disease [34-36] heralding an investigation of its utility for motivational deficits in HIV-1.

Thus, the goals of the present study were twofold. First, to investigate goal-directed behavior, an index of apathy, in older (i.e., 7-9 months of age) HIV-1 transgenic (Tg) rats. Assessments of apathy in young (i.e., approximately 2 months of age) HIV-1 Tg rats revealed apathetic behavior characterized by decreased response vigor and diminished sensitivity to, and reinforcing efficacy of, cocaine [37]. Second, to systematically evaluate the therapeutic utility of SE for apathy in older HIV-1 Tg rats. Given that pharmacological treatments for apathy associated with neurodegenerative diseases are currently limited [38], the development of a novel therapeutic for apathy associated with HIV-1 has the potential for broad clinical significance.

## METHODS

### Animals

Motivational alterations associated with HIV-1, and the efficacy of SE as an innovative therapeutic, were evaluated in ovariectomized (OVX) female Fisher (F344/NHsd; Harlan Laboratories, Inc., Indianapolis, IN) HIV-1 Tg (*n* = 21) and control (*n* = 21) rats. Resembling HIV-1 seropositive individuals on cART, the HIV-1 Tg rat, originally reported by Reid et al. [39], expresses HIV-1 viral proteins constitutively throughout development [40-41]. The HIV-1 Tg rat exhibits relatively good health throughout the functional lifespan, evidenced by similar growth rates relative to control animals [25, 40, 42] and intact sensory and motor system function [40, 43].

Unrelated animals were requested to prevent the violation of the independent observation assumption inherent in many traditional statistical techniques. Furthermore, given the potential for sporadic transgene insertion elsewhere in the rat genome, age-matched F344/NHsd controls (rather than littermates) were purchased from Harlan Laboratories to assure the most developmentally appropriate and genetically stable baseline.

Animals were delivered to the animal vivarium, after being OVX at Harlan Laboratories, between six and eight months of age. To preclude any potential confounding effects of endogenous hormones, animals were OVX and fed a minimal phytoestrogen diet (≤20 ppm; Teklad 2020X Global Extruded Rodent Diet (Soy Protein-Free)). Animals had *ad libitum* access to rodent food and water, unless otherwise specified.

Animals were maintained in AAALAC-accredited facilities according to guidelines established by the National Institutes of Health. The animal colony was maintained at 21^°^ ± 2^°^C, 50±10% relative humidity and a 12-h light:12-h dark cycle with lights on at 0700h (EST). The Institutional Animal Care and Use Committee (IACUC) at the University of South Carolina (Federal Assurance #D16-00028) approved all experimental procedures.

### Apparatus

Assessments of goal-directed behavior were conducted in operant chambers (ENV-008; Med-Associates, St. Albans, VT, USA) located within sound-attenuating enclosures and controlled by Med-PC computer interface software. The front panel of the operant chamber contained a magazine that allowed a recessed 0.01cc dipper cup (ENV-202C) to deliver a solution through a 5cm x 5cm opening (ENV 202M-S), two retractable “active” metal levers (i.e., responding resulted in reinforcement; ENV-112BM), and two white cue lights (28 volts). Head entries into the magazine were detected using an infrared sensor (ENV 254-CB). To control for side bias, an “active” lever retracted after five consecutive presses on a single lever. The rear panel of the operant chamber had one non-retractable “inactive” lever, whereby responding was recorded, but not reinforced, and a house light (28 volts).

During Cocaine-Maintained Responding, intravenous cocaine infusions were delivered through a water-tight swivel (Instech 375/22ps 22GA; Instech Laboratories, Inc., Plymouth Meeting, PA), which was connected to the backmount of the animal using Tygon tubing (ID, 0.020 IN; OD, 0.060 IN) enclosed by a stainless steel tether (Camcaths, Cambridgeshire, Great Britain), using a syringe pump (PHM-100). A Med-PC computer program was utilized to calculate pump infusion times based on an animal’s daily bodyweight.

### Drugs

Cocaine hydrochloride (Sigma-Aldrich Pharmaceuticals, St. Louis, MO) was weighed as a salt and was dissolved in physiological saline (0.9%; Hospira, Inc., Lake Forest, IL). To preclude any significant hydrolysis of cocaine [44], solutions were prepared prior to the start of each testing session for every animal. Sucrose solutions were prepared at the beginning of each testing day. After obtaining SE from Cayman Chemical Company (Ann Arbor, MI, USA), it was incorporated into 100 mg sucrose pellets (0.05 mg SE per sucrose pellet) by Bio-Serv (Flemington, NJ). 100 mg sucrose pellets were purchased from Bio-Serv for the vehicle group.

### Experimental Timeline

The experimental timeline for SE treatment the series of three experiments evaluating goal-directed behavior, and neuroanatomical assessments is illustrated in Figure 1.

**FIGURE 1.**
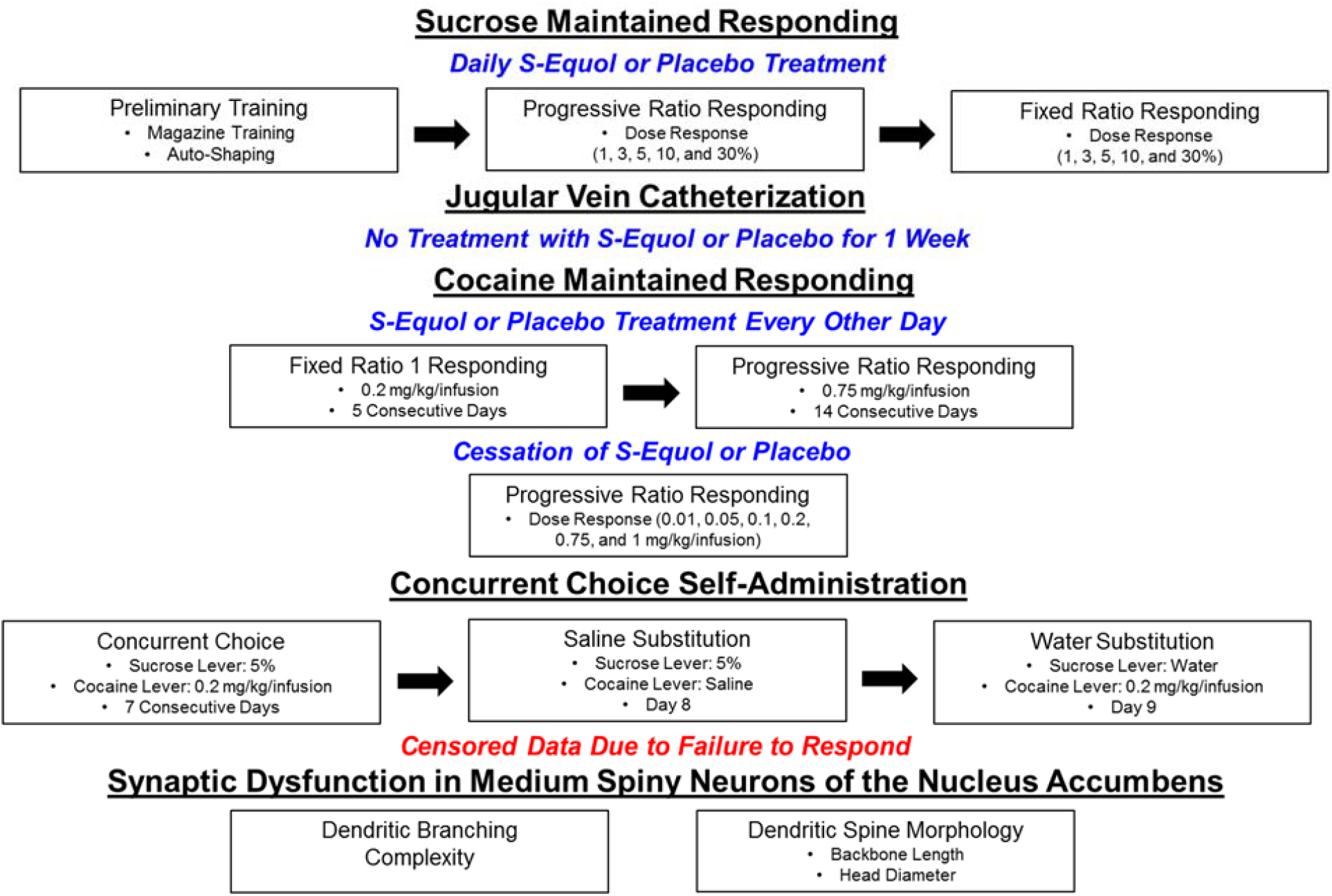
Experimental Design Schematic.

### Treatment

Animals were randomly assigned to receive either 0.2 mg SE (Control: *n*=11; HIV-1 Tg: *n*=11) or vehicle (Control: *n*=10; HIV-1 Tg: *n*=10). Between approximately 7 and 9 months of age, and one week prior to the start of operant training, HIV-1 Tg and control animals began receiving a daily treatment with either SE or vehicle. Given that each sucrose pellet contained 0.05 mg SE, animals receiving SE treatment received four pellets per day. Animals receiving vehicle treatment received four sucrose pellets per day. Daily treatment continued until jugular vein catheterization. Following jugular vein catheterization, animals were not treated for one week. When treatment resumed, it was given every other day until the end of the 14-day cocaine self-administration PR task. No treatment occurred during the cocaine self-administration PR dose-response task or concurrent choice self-administration. In total, HIV-1 Tg and control animals were treated for 84 days.

The 0.2 mg dose of SE was selected for two complementary reasons. First, using a dose-response experiment design, 0.2 mg SE was established as the most efficacious dose for the alleviation of sustained attention deficits in the HIV-1 Tg rat [34]. Second, the dose selected yielded a daily amount of 0.25-1.0 mg/kg SE (i.e., equivalent to a 2.5-10 mg dose in a 60 kg human); a dose well below the daily isoflavone intake of most elderly Japanese (i.e., 30-50 mg; [45]).

### Sucrose Maintained Responding

#### Preliminary Training

Previously established research protocols were used to conduct dipper training and autoshaping [37, 46-48]. During dipper training, animals were trained to approach the magazine and drink a 5% sucrose solution (w/v) from the dipper receptacle. Subsequently, during autoshaping, animals learned to lever press for the 5% sucrose solution (w/v) using a fixed-ratio (FR) 1 schedule of reinforcement. Water restriction (12-15 hours prior to assessment) was implemented throughout preliminary training. Animals had *ad libitum* access to water for 9-12 hours after the completion of testing.

To successfully acquire autoshaping, animals were required to achieve at least 60 reinforcers for 3 consecutive days. *Ad libitum* access to water was reinstated following the successful completion of preliminary training.

#### Progressive Ratio Responding: Dose Response

Following the completion of preliminary training, animals completed one maintenance session (i.e., 5% sucrose on an FR-1 schedule of reinforcement) after which animals were assessed using a progressive ratio (PR) schedule of reinforcement (Maximum Session Length: 120 Minutes). A dose-response experimental design was utilized, whereby the reinforcer was one of five sucrose concentrations (1, 3, 5, 10, and 30% w/v), presented using a Latin-Square experimental design, on test days, which occurred every other day. A maintenance session also occurred on intervening non-test days.

During PR tests, the ratio requirement was completed by responding across the two “active” levers. After successfully meeting the ratio requirement, the active levers were retracted and animals had 4 seconds of access to sucrose. The ratio requirement (rounded to the nearest integer) increased progressively according to the following exponential function: [5e^(reinforcer number X 0.2)^]-5 [49].

#### Fixed Ratio Responding: Dose Response

Subsequently, animals responded for the same dose-response sucrose concentrations (i.e., 1, 3, 5, 10, and 30% w/v) on an FR-1 schedule of reinforcement. Testing days occurred every other day and a maintenance session occurred on the intervening non-test days.

### Cocaine Maintained Responding

#### Jugular Vein Catheterization

Following the completion of sucrose maintained responding, jugular vein catheterization was performed using the methods reported by Bertrand et al. [37]. In brief, HIV-1 Tg and control animals were anesthetized using 5% inhalant sevofluorane and maintained at 3.5-4% sevofluorane throughout the surgical procedure. After anesthesia induction, a sterile IV catheter, which extended dorsally and connected to an acrylic pedestal embedded with mesh, was implanted into the right jugular vein and secured with sterile sutures (4-0 Perma-Hand silk; EthiconEnd-Surgery, Inc.). The dorsal portion of the catheter/backmount was implanted subcutaneously above the right and left scapulae and stitched into place using sterile, absorbable sutures (4-0 Monoweb). Immediately following surgery, post-operative analgesia was provided by butorphenol (Dorolex; 0.8 mg/kg, SC; Merck Animal Health, Millsboro, DE) and the antibiotic gentamicin sulfate (0.2 ml 1%, IV; VEDCO, Saint Joseph, MO) was administered to prevent infection. Rats were monitored in a heat-regulated warm chamber following surgery and returned to the colony room after recovery from anesthesia. Two HIV-1 Tg animals died immediately following surgery; one of unknown causes and one suffered from a seizure after returning to the home cage, yielding Control Vehicle, *n*=10; Control SE, *n*=11; HIV-1 Tg Vehicle, *n*=9; HIV-1 Tg SE, *n*=10 for Cocaine-Maintained Responding.

For one week following surgery, catheters were “flushed” daily with a solution containing heparin (2.5%; APP Pharmaceuticals, Schamburg, IL) and the antibiotic gentamicin sulfate (1%) to prevent clotting and infections. Seven days after jugular vein catheterization, animals began Cocaine-Maintained Responding and resumed treatment with either SE or vehicle. Prior to operant testing each day, catheters were “flushed” with 0.9% saline solution (Baxter, Deerfield, IL). After the completion of daily operant testing, catheters were “flushed” with post-flush solution.

#### Response Escalation

Two phases (i.e., FR-1 Responding and PR Responding), modified from Morgan et al. [50] and previously employed in our laboratory by Bertrand et al. [37], were utilized to produce an escalation of cocaine-maintained responding. First, rats responded for cocaine (0.2 mg/kg/inf) according to a FR-1 schedule of reinforcement for 5 consecutive days (Session Length: 1 hour). Second, rats responded for cocaine (0.75 mg/kg/inf) on a PR schedule of reinforcement, whereby the ratio requirement was defined using the exponential function defined above for 14 consecutive days (Maximum Session Length: 120 Minutes). During both FR-1 and PR responding, a 20s time-out (i.e., active levers were retracted and the house light was extinguished) following the completion of a response requirement.

#### Progressive Ratio Responding: Dose Response

Subsequently, a dose response experimental design was utilized to evaluate the reinforcing efficacy of cocaine. Cocaine concentrations (0.01, 0.05, 0.1, 0.2, 0.75, and 1.0 mg/kg/inf) were presented in ascending order. A maintenance session (i.e., 0.2 mg/kg/ inf on an FR-1 schedule of reinforcement) occurred every other day.

### Concurrent Choice Self-Administration

After establishing a history of sucrose and cocaine maintained-responding, choice behavior was evaluated using a concurrent choice self-administration experimental paradigm for seven consecutive days. Throughout the experimental procedure, animals were responding for 5% (w/v) sucrose or 0.2 mg/kg/inf of cocaine. On the eighth day, saline was substituted for cocaine. On the ninth day, water was substituted for sucrose. Sucrose-paired lever presentation (right or left) was balanced between groups.

Each session began with four forced choice trials (i.e., only one “active” lever was available) where animals responded for two cocaine and two sucrose reinforcers. Following the forced choice trials, both “active” levers were concurrently available to allow the animals to freely choose between sucrose and cocaine. After a response was made, a 20s time-out occurred.

Data for the concurrent choice self-administration were censored due to the absence of responding, and thus absence of a choice, independent of genotype group. Censoring reduced group sample sizes by approximately 50%. Statistical power estimates for a three-way interaction (Genotype x Reinforcer x Day) investigating the impact of the HIV-1 transgene on choice behavior was 0.31; an estimate which is significantly lower than appropriate statistical power levels (i.e., 0.8; [51]) and thus precluded drawing any reliable inferences about the choice experiment.

### Synaptic Dysfunction in Medium Spiny Neurons of the Nucleus Accumbens

#### Preparation of Tissue

Animals were transcardially perfused within 24 hours of their last self-administration session. After animals were deeply anesthetized using sevoflurane (Abbot Laboratories, North Chicaco, IL), transcardial perfusion was conducted using the methodology reported by Roscoe et al. [25] with one minor modification. Specifically, after the brains were dissected, they were post-fixed in 4% paraformaldehyde for 10 minutes.

#### DiOListic Labeling

MSNs from the NAc were visualized using a ballistic labeling technique, originally described by Seabold et al. [52]. Methodology for the preparation of DiOlistic cartridges and Tefzel tubing and DiOlistic labeling was previously described in detail [25].

#### Dendritic Analysis and Spine Quantification: Medium Spiny Neurons

MSNs were analyzed from the NAc, located approximately 2.28 mm to 0.60 mm anterior to Bregma [53]. Z-stack images were obtained using methodology previously reported [25].

Dendritic branching was manually evaluated for images with a clear dendritic arbor, as assessed by the maximum intensity projection image. Analyses were conducted on one to four MSNs per animal, yielding Control Vehicle, *n*=7; Control SE, *n*=7; HIV-1 Tg Vehicle, *n*=5; HIV-1 Tg SE, *n*=10. Dendritic spine parameters were analyzed using the AutoNeuron and AutoSpine extension modules in Neurolucida360 (MicroBrightfield, Williston, VT, USA). Spine analyses were conducted for MSNs meeting the following selection criteria, including 1. Continuous staining beginning in the cell body extending throughout the dendrite; 2. Minimal DiI diffusion; and 3. Low background fluorescence. Analyses were conducted on one to eight MSNs per animal, yielding Control Vehicle, *n*=10; Control SE, *n*=11; HIV-1 Tg Vehicle, *n*=8; HIV-1 Tg SE, *n*=10. Given the nested experimental design, cluster means were calculated for both dendritic branching and spine parameters, whereby the sample size (*n*) reflects the number of animals in each group.

#### Dendritic Spine Morphology

Two dendritic spine morphological parameters, including dendritic spine backbone length (µm) and dendritic spine head diameter (µm), were assessed. Dendritic spines were included in the analysis if they met the definitional criteria established using previously published manuscripts (i.e., backbone length, 0.2 to 4.0 µm [54]; head diameter, 0.0 to 1.2 µm [55]; volume, 0.05 to 0.85 µm^3^ [56]).

### Statistical Analysis

Data were analyzed using analysis of variance (ANOVA) and regression techniques (SAS/STAT Software 9.4, SAS Institute, Inc., Cary, NC; SPSS Statistics 27, IBM Corp., Somer, NY; GraphPad Software, Inc., La Jolla, CA). An alpha criterion of p≤0.05 was utilized to establish statistical significance. Orthogonal decompositions and/or the Greenhouse-Geisser df correction factor [57] were implemented to account for variables that may violate the compound symmetry assumption. Based on the *a priori* aims of the present study, planned comparisons were conducted to evaluate the impact of chronic HIV-1 viral protein exposure on goal-directed behavior (i.e., Control Vehicle vs. HIV-1 Tg Vehicle), the presence of an SE effect (i.e., HIV-1 Tg Vehicle vs. HIV-1 Tg SE), and the magnitude of the SE effect (i.e., Control Vehicle vs. HIV-1 Tg SE).

For sucrose-maintained responding, the *a priori* planned comparisons were addressed using a regression and/or ANOVA techniques. Specifically, regression analyses were conducted to evaluate the shape and parameters of the best-fit function for the acquisition of autoshaping, PR responding, and FR-1 responding. To further evaluate the impact of chronic HIV-1 viral protein exposure on PR responding, a mixed-model ANOVA was conducted using SPSS Statistics 27. Two dependent variables of interest, including the number of reinforcers and the number of “active” lever presses, were investigated. In cases where an animal was tested more than once at a single concentration, a cluster mean was calculated to account for the nested experimental design [58-59]. The mean series imputation method was used for all censored data, which occurred at the 1% (HIV-1 Tg SE: *n*=1) and 10% (Control Vehicle: *n*=1) concentration. One HIV-1 Tg animal treated with vehicle failed to acquire autoshaping and therefore did not complete either the PR or FR assessment; the animal was not included in the statistical analyses for these tasks.

For cocaine-maintained responding, regression and/or ANOVA techniques were also utilized to statistically evaluate the *a priori* planned comparisons. FR-1 responding for a 0.2 mg/kg/inf dose of cocaine was evaluated using regression analyses and a mixed-model ANOVA (PROC MIXED; SAS/STAT Software 9.4) with an unstructured covariance structure. The number of infusions earned and number of “active” lever presses during the cocaine PR task (0.75 mg/kg/inf dose) were evaluated using regression analyses. Additionally, a mixed model ANOVA (PROC MIXED; SAS/STAT Software 9.4) with a compound symmetry covariance structure was also conducted to assess the number of “active” lever presses during the cocaine PR task. Regression analyses were used to statistically analyze the number of cocaine infusions during the dose-response cocaine PR assessment.

Neuronal morphology and dendritic spine morphological parameters were analyzed using ANOVA techniques. Specifically, dendritic spine branch order was evaluated using a mixed-model ANOVA (SPSS Statistics 27) and dendritic spine backbone length and head diameter were analyzed using a generalized linear mixed effects model with a Poisson distribution and an unstructured covariance structure (PROC GLIMMIX; SAS/STAT Software 9.4).

## Data Availability Statement

All relevant data are within the paper.

## RESULTS

### Presence of the HIV-1 Transgene

#### Sucrose Maintained Responding

##### Acquisition of Autoshaping

At the genotypic level, HIV-1 Tg animals treated with vehicle acquired autoshaping, earning at least 60 reinforcers for three consecutive days, significantly slower than control animals treated with vehicle (Figure 2A). After 66 days, 100% of the control and 90% of the HIV-1 Tg rats treated with vehicle met criteria. A sigmoidal-dose response curve (variable slope) afforded the best-fit for the number of days to criteria for both HIV-1 Tg (*R*^*2*^= 0.98) and control (*R*^*2*^= 0.98) animals treated with vehicle; albeit statistically significant differences in the parameters of the function [*F*(4, 27)=4.6, *p≤*0.006] were observed. Results support, therefore, a prominent deficit in stimulus-reinforcement learning in HIV-1 Tg rats treated with vehicle.

**FIGURE 2.**
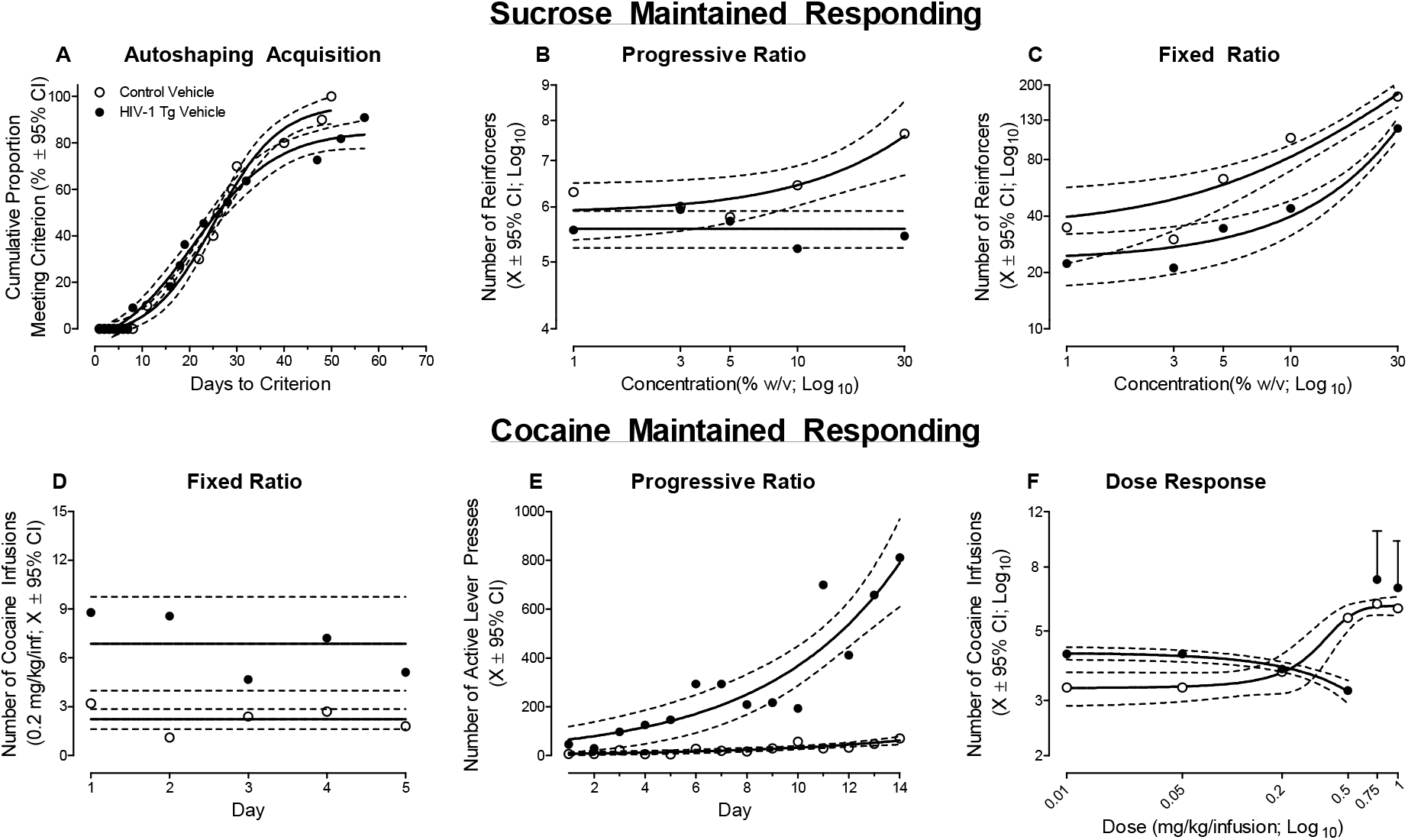
The impact of the HIV-1 transgene on sucrose-maintained responding (**A-C**) and cocaine-maintained responding (**D-F**) is illustrated as a function of genotype (HIV-1 Tg Vehicle vs. Control Vehicle; ±95% Confidence Intervals). HIV-1 Tg animals took significantly longer to successfully acquire autoshaping, supporting a profound deficit in stimulus-reinforcement learning (**A**). Under sucrose-maintained responding, apathetic behaviors in the HIV-1 Tg rat were characterized by a diminished sensitivity to (**B**), and reinforcing efficacy of (**C**), sucrose relative to control animals. Under cocaine-maintained responding, HIV-1 Tg animals exhibited an increased response vigor, independent of schedule of reinforcement (i.e., Fixed Ratio 1: **D**; Progressive Ratio: **E**), relative to control animals. Furthermore, an increased sensitivity to cocaine dose was observed in HIV-1 Tg animals relative to controls (**F**). Collectively, independent of differences in the unique response to either sucrose or cocaine self-administration, the HIV-1 Tg rat exhibits prominent alterations in goal-directed behaviors.

##### Progressive Ratio Responding: Dose Response

After successfully acquiring autoshaping, the reinforcing efficacy of sucrose was evaluated using a dose-response experimental design and PR schedule of reinforcement. HIV-1 Tg animals treated with vehicle exhibited a diminished reinforcing efficacy of sucrose relative to control animals treated with vehicle evidenced by a statistically significant genotype x concentration interaction for both the number of reinforcers (Figure 2B; [*F*(4, 68)=3.4, *p*_GG_≤0.02, η^2^ _p_ =0.165] with a prominent linear component [*F*(1,17)=6.7, *p*≤0.02, η^2^ _p_ =0.282]) and “active” lever presses (Data Not Shown; [*F*(4, 68)=2.9, *p*_GG_≤0.05, η^2^ _p_ =0.147] with a prominent linear component [*F*(1,17)=6.0, *p*≤0.03, η^2^ _p_ =0.261]. Specifically, control animals treated with vehicle displayed a linear increase in the number of reinforcers (*R*^*2*^= 0.87) and “active” lever presses (*R*^*2*^= 0.93) as sucrose concentration increased. The number of reinforcers earned and “active” lever presses produced by HIV-1 Tg animals treated with vehicle, however, remained invariant across sucrose concentration, well-described by a horizontal line. Older HIV-1 Tg animals, therefore, exhibit a diminished reinforcing efficacy of sucrose.

##### Fixed Ratio Responding: Dose Response

The sensitivity to sucrose concentration was subsequently examined using an FR-1 schedule of reinforcement. HIV-1 Tg animals treated with vehicle exhibited a blunted sensitivity to sucrose concentration relative to control animals treated with vehicle (Figure 2C), evidenced by a differential pattern of responding. Regression analyses revealed a dose-response function well-characterized by a one-phase association in control animals treated with vehicle (*R*^*2*^= 0.95), whereas an exponential growth equation afforded a best-fit for HIV-1 Tg animals treated with vehicle (*R*^*2*^= 0.98).

Collectively, results support a prominent deficit in stimulus-reinforcement learning, as well as apathetic behaviors in older HIV-1 Tg rats; apathetic behaviors that are characterized by a diminished reinforcing efficacy of, and sensitivity to, sucrose.

#### Cocaine Maintained Responding

##### Fixed Ratio Responding

Following jugular vein catheterization, HIV-1 Tg and control animals responded for 0.2 mg/kg/inf of cocaine according to a FR-1 schedule of reinforcement for 5 consecutive days. HIV-1 Tg animals treated with vehicle displayed an increased response vigor, independent of day, relative to control animals treated with vehicle (Figure 2D; Main Effect: Genotype, [*F*(1,17)=4.5, *p*≤0.05]). Furthermore, the number of cocaine infusions across day was well-described by a horizontal line for both HIV-1 Tg and control animals treated with vehicle; albeit statistically significant differences in the mean of the function were observed [*F*(1,93)=11.1, *p*≤0.01]). Thus, HIV-1 Tg animals treated with vehicle exhibited increased response vigor for cocaine, supporting enhanced drug seeking behavior, relative to control rats treated with vehicle. *Progressive Ratio Responding*. Independent of genotype, both HIV-1 Tg and control animals treated with vehicle escalated their drug intake, evidenced by an increase in the number of “active” lever presses (Figure 2E) and the number of cocaine infusions (Data Not Shown; 0.75 mg/kg/inf), across the 14 consecutive days of PR testing.

With regards to the number of “active” lever presses, HIV-1 Tg animals treated with vehicle escalated responding at a significantly faster rate relative to control animals treated with vehicle (Day x Genotype interaction, [*F*(1, 245)=12.0, *p*≤0.01]). An exponential growth equation afforded the best fit for the number of “active” lever presses for both HIV-1 Tg (*R*^*2*^= 0.82) and control (*R*^*2*^= 0.73) animals treated with vehicle; albeit statistically significant differences in the parameters of the function were observation [*F*(2,24)=66.5, *p*≤0.01].

Collectively, results support enhanced drug seeking in HIV-1 Tg animals, evidenced by increase response vigor and faster escalation, relative to control animals treated with vehicle; an observation that is independent of schedule of reinforcement (i.e., FR vs. PR).

#### Progressive Ratio Responding: Dose Response

HIV-1 Tg animals treated with vehicle exhibited an increased reinforcing efficacy of cocaine dose (mg/kg/inf) relative to control animals treated with vehicle (Figure 2F). Specifically, in HIV-1 Tg animals treated with vehicle, a linear (*R*^*2*^= 0.98) decrease in the number of cocaine infusions was observed from the 0.01 to 0.5 mg/kg/inf dose; a dramatic increase in the number of infusions was observed at the 0.75 mg/kg/inf and 1 mg/kg/inf dose. In sharp contrast, in control animals treated with vehicle, the number of cocaine infusions across all doses was well-described by a sigmoidal-dose response curve (variable slope; *R*^*2*^= 0.99). Collectively, results support enhanced drug seeking behavior and an increased differential reinforcing efficacy of cocaine in older HIV-1 Tg rats treated with vehicle relative to control rats treated with vehicle.

#### Synaptic Dysfunction in Medium Spiny Neurons of the Nucleus Accumbens

##### Branch Order

Dendritic branching was evaluated by counting the number of primary, secondary, and tertiary branches in MSNs of the NAc. Following chronic HIV-1 viral protein exposure, a profound decrease in dendritic branching complexity was observed in HIV-1 Tg animals treated with vehicle relative to control animals treated with vehicle (Figure 3A; Genotype x Branch Order interaction with a prominent linear component, [*F*(1,10)=7.3, *p*≤0.02, η^2^ _p_=*0*.422]). Thus, presence of the HIV-1 transgene leads to prominent decreases in dendritic branching complexity in MSNs of the NAc.

**FIGURE 3.**
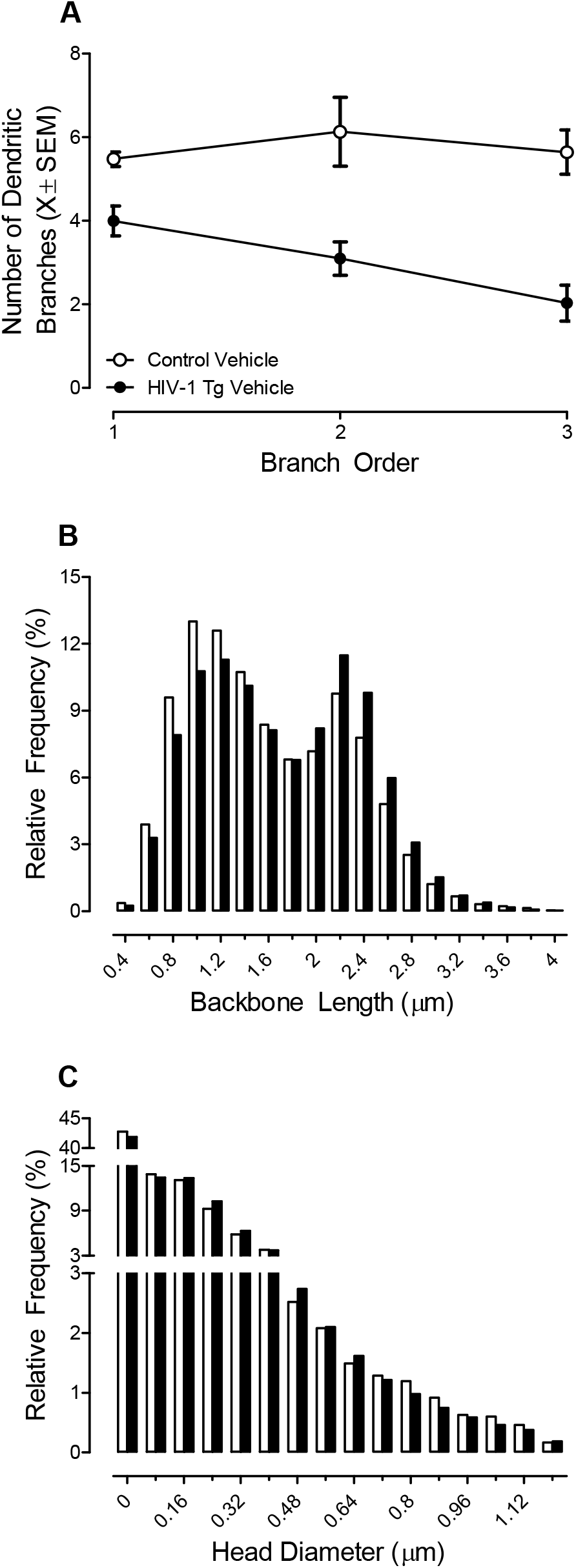
Following a history of cocaine self-administration, the impact of the HIV-1 transgene on dendritic branching (**A**; ±SEM) and dendritic spine morphology (**B-C**) in medium spiny neurons of the nucleus accumbens was examined and is illustrated as a function of genotype (HIV-1 Tg Vehicle vs. Control Vehicle). HIV-1 Tg animals exhibited a dramatic decrease in dendritic branching complexity (**A**) and a population shift towards longer dendritic spines (**B**) with decreased head diameter (**C**) relative to control animals.

##### Dendritic Spine Morphology

After a history of cocaine self-administration, dendritic spine morphology was examined in MSNs of the NAc. HIV-1 Tg animals treated with vehicle exhibited a prominent population shift towards longer dendritic spines (Figure 3B; [*F*(1,1424)=386.2, *p*≤0.01]) with decreased head diameter (Figure 3C; [*F*(1,1196)=16.8, *p*≤0.01]) relative to control animals treated with vehicle. Presence of the HIV-1 transgene, therefore, shifts the morphological parameters of dendritic spines in MSNs of the NAc to a more immature phenotype.

### Therapeutic Efficacy of S-Equol

#### Sucrose Maintained Responding

##### Acquisition of Autoshaping

Treatment with SE ameliorated deficits in stimulus-reinforcement learning in HIV-1 Tg animals. All HIV-1 Tg animals treated with SE successfully acquired autoshaping. The best-fit function (i.e., global sigmoidal dose-response curve (variable slope)) for the number of days to criteria for HIV-1 Tg animals treated with SE was statistically indistinguishable from either HIV-1 Tg animals treated with vehicle (Figure 4A; *p*>0.05; *R*^*2*^= 0.98) or control animals treated with vehicle (Figure 4B; *p*>0.05; *R*^*2*^= 0.98). Thus, the marked impairment in stimulus-reinforcement learning observed in HIV-1 Tg animals treated with vehicle was mitigated by treatment with SE.

**FIGURE 4.**
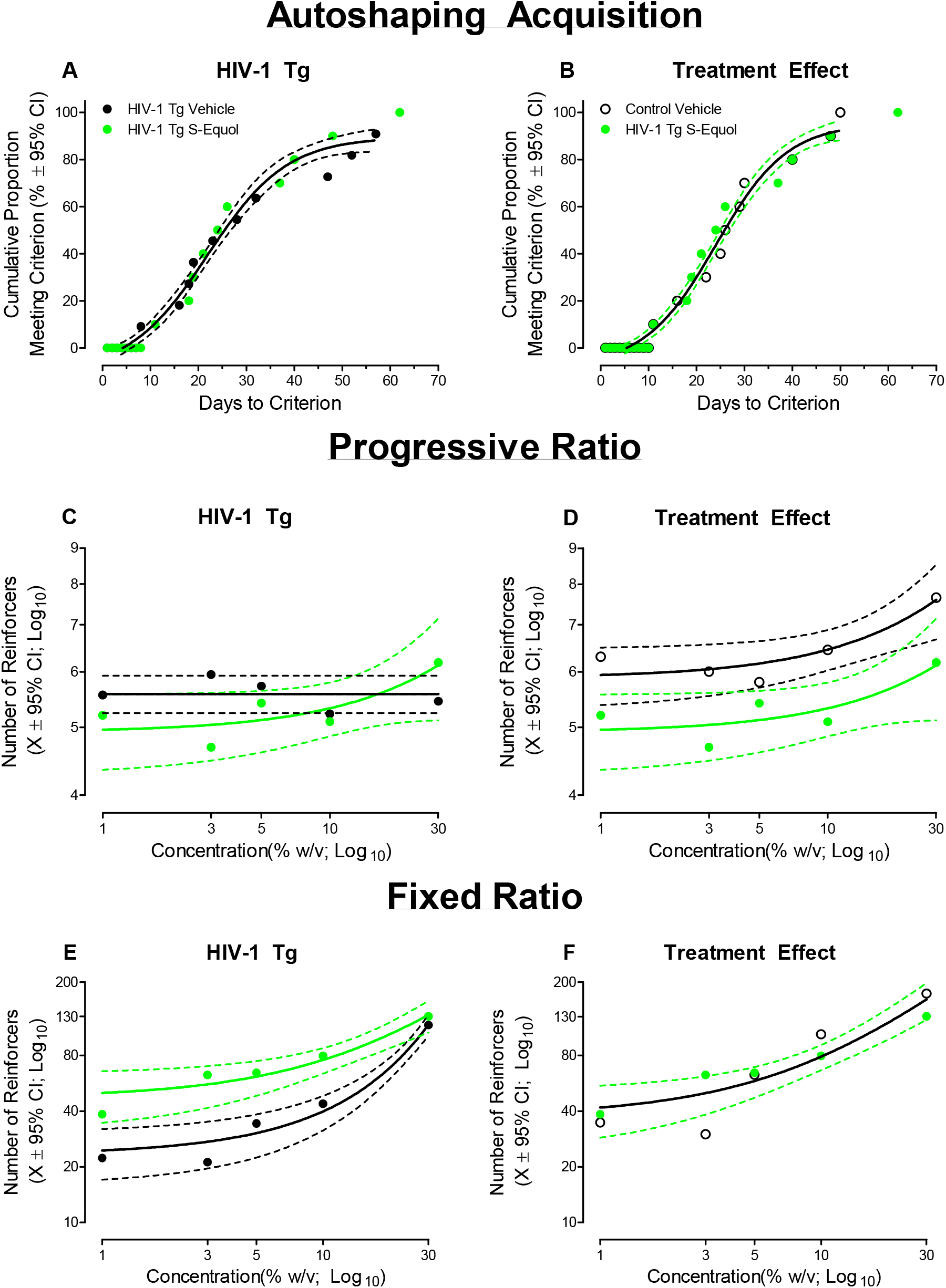
The therapeutic efficacy of S-Equol (SE) as a novel therapeutic for apathetic behaviors under sucrose-maintained responding in the HIV-1 Tg rat is illustrated as a function of genotype (HIV-1 Tg vs. Control) and treatment (SE vs. Vehicle; ±95% Confidence Intervals). The number of days required to successfully acquire autoshaping, an index of stimulus-reinforcement learning, for HIV-1 Tg animals treated with SE was statistically indistinguishable from either HIV-1 Tg animals treated with vehicle (**A**) or control animals treated with vehicle (**B**). Treatment with SE enhanced the reinforcing efficacy of sucrose (**C-D**) and ameliorated alterations in the sensitivity to sucrose (**E-F**) in HIV-1 Tg rats.

##### Progressive Ratio Responding: Dose Response

The reinforcing efficacy of sucrose in HIV-1 Tg animals was enhanced by treatment with SE. First, comparison of HIV-1 Tg animals treated with SE and HIV-1 Tg animals treated with vehicle revealed a differential pattern of responding. Specifically, HIV-1 Tg animals treated with SE exhibited a linear increase in the number of reinforcers (Figure 4C; *R*^*2*^= 0.73) and “active” lever presses (Data Not Shown; *R*^*2*^= 0.91) as sucrose concentration increased; a concentration-dependent effect not observed in HIV-1 Tg animals treated with placebo (Best Fit: Horizontal Line).

Second, both HIV-1 Tg animals treated with SE and control animals treated with vehicle exhibited a linear increase in the number of reinforcers (Figure 4D; *R*^*2*^= 0.73 and 0.87, respectively) and “active” lever presses (Data Not Shown; *R*^*2*^= 0.93 and 0.91, respectively) as sucrose concentration increased. Although HIV-1 Tg animals treated with SE exhibited decreased response vigor (i.e., Statistically Significant Difference in β_0_; Reinforcers: [*F*(1,6)=12.4, *p*≤0.01]; “Active” Lever Presses: [*F*(1,6)=10.8, *p*≤0.02]) relative to control animals treated with vehicle, the rate of increase (i.e., β_1_) across sucrose concentration was statistically indistinguishable between groups (*p*>0.05). Treatment with SE, therefore, mitigates the diminished reinforcing efficacy of sucrose observed in older HIV-1 Tg animals.

##### Fixed Ratio Responding: Dose Response

Alterations in the sensitivity to sucrose were ameliorated by treatment with SE in HIV-1 Tg animals. Comparison of HIV-1 Tg animals treated with SE and HIV-1 Tg animals treated with vehicle (Figure 4E) revealed differential patterns of responding (i.e., First-Order Polynomial (*R*^*2*^= 0.95) and Exponential Growth Equation (*R*^*2*^= 0.98), respectively), whereby HIV-1 Tg animals treated with SE exhibited greater sensitivity to varying sucrose concentrations. Furthermore, the number of sucrose reinforcers earned by HIV-1 Tg animals treated with SE and control animals treated with vehicle (Figure 4F) was well-described by a global one-phase association (*R*^*2*^= 0.91) supporting no statistically significant differences between groups in sucrose sensitivity. Taken together, treatment with SE mitigated deficits in stimulus-reinforcement learning and apathetic behaviors in older HIV-1 Tg rats by enhancing the reinforcing efficacy of, and sensitivity to, sucrose.

#### Cocaine Maintained Responding

##### Fixed Ratio Responding

HIV-1 Tg animals treated with SE displayed an initial novelty response to cocaine self-administration followed by a rapid decay, to levels statistically indistinguishable from either HIV-1 Tg or control animals treated with vehicle, in the number of cocaine infusions earned across the five day testing period.

Comparison of HIV-1 Tg animals treated with SE and HIV-1 Tg animals treated with vehicle revealed a differential pattern of responding (Figure 5A; Treatment x Day interaction, [*F*(1,74)=4.42, *p*≤0.04]). Specifically, a one-phase decay afforded the best-fit for the number of infusions earned by HIV-1 Tg animals treated with SE (*R*^*2*^= 0.97), while a horizontal line was most appropriate for HIV-1 Tg animals treated with placebo. However, the overlapping 95% confidence intervals observed from self-administration days two through five suggest that the differential pattern of responding is driven primarily by an initial novelty response in HIV-1 Tg animals treated with SE on the first day of cocaine self-administration.

**FIGURE 5.**
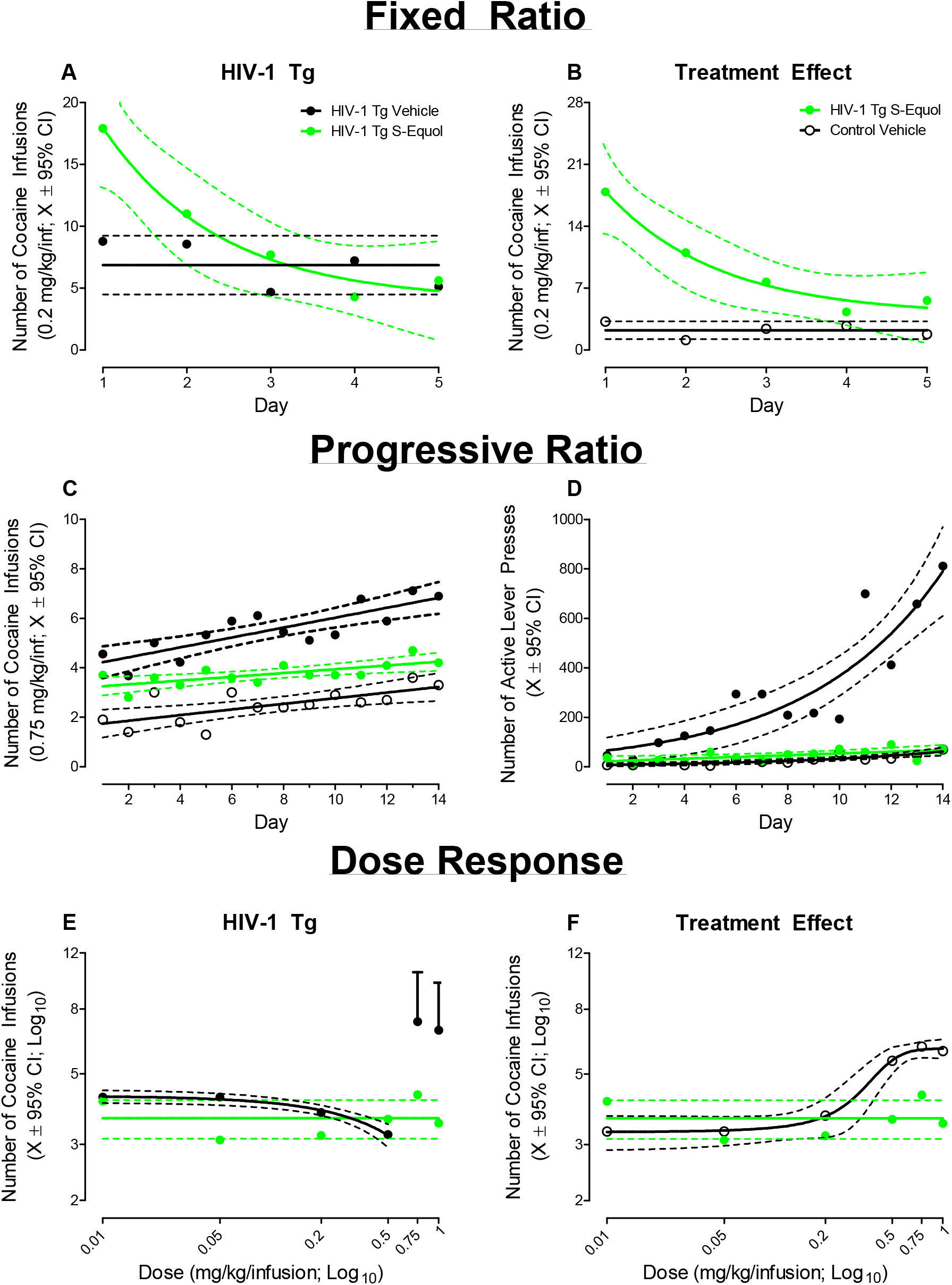
The therapeutic efficacy of S-Equol (SE) as a novel therapeutic for apathetic behaviors under cocaine-maintained responding in the HIV-1 Tg rat is illustrated as a function of genotype (HIV-1 Tg vs. Control) and treatment (SE vs. Vehicle; ±95% Confidence Intervals). HIV-1 Tg animals treated with SE displayed an initial novelty response to cocaine self-administration followed by a rapid decay. At the end of the five day fixed ratio testing period, the number of cocaine infusions earned by HIV-1 Tg animals treated with SE were statistically indistinguishable from either HIV-1 Tg (**A**) or control animals (**B**) treated with vehicle. Treatment with SE reduced drug seeking behavior in the HIV-1 Tg rat (**C-D**); an effect which generalized across cocaine dose (**E-F**).

A differential pattern of responding (i.e., HIV-1 Tg SE: One-Phase Decay, *R*^*2*^= 0.97; Control Vehicle: Horizontal Line) was also observed when comparing HIV-1 Tg animals treated with SE and control animals treated with vehicle (Figure 5B; Genotype x Day interaction, [*F*(1,78)=12.4, *p*≤0.01]). Again, however, the overlapping 95% confidence intervals observed at self-administration days four and five suggests that the number of cocaine infusions earned by HIV-1 Tg animals treated with SE at the end of FR-1 responding is statistically indistinguishable from control animals treated with vehicle.

##### Progressive Ratio Responding

Independent of genotype and/or treatment, an escalation of cocaine intake, evidenced by an increase in the number of cocaine infusions (Figure 5C) and number of “active” lever presses (Figure 5D), was observed across the 14 consecutive days of PR testing.

Examination of the number of cocaine infusions revealed a linear increase in the number of cocaine infusions for all three groups (Control Vehicle: *R*^*2*^= 0.48, HIV-1 Tg Vehicle: *R*^*2*^= 0.69, HIV-1 Tg SE: *R*^*2*^= 0.49). Treatment with SE, however, mitigated response vigor and drug escalation relative to HIV-1 Tg animals treated with vehicle. Specifically, comparison of HIV-1 Tg animals treated with SE and HIV-1 Tg animals treated with vehicle revealed statistically significant differences in the intercept (i.e., β_0_; [*F*(1, 24)=5.0, *p*≤0.04]) and in the rate of escalation (i.e., β_1_; [*F*(1,24)=7.8, *p*≤0.01]) between groups. Albeit, response vigor in HIV-1 Tg animals treated with SE was still significantly greater than control animals treated with vehicle (β_0_: [*F*(1,24)=30.0, *p*≤0.01]; no statistically significant differences in the rate of escalation (i.e., β_1_) were observed (*p*>0.05).

With regards to the number of “active” lever presses, HIV-1 Tg animals treated with SE escalated responding significantly slower than HIV-1 Tg animals treated with vehicle (Day x Treatment interaction, [*F*(1,245)=12.2, *p*≤0.01]). Specifically, HIV-1 Tg animals treated with SE and HIV-1 Tg animals treated with vehicle exhibited differential patterns of escalation, evidenced by different best-fit functions (i.e., First-Order Polynomial, *R*^*2*^= 0.39 and Exponential Growth Equation, *R*^*2*^= 0.82, respectively). Critically, the escalation of responding in HIV-1 Tg animals treated with SE was statistically indistinguishable from control animals treated with vehicle (Day x Treatment interaction, *p*>0.05). Thus, treatment with SE reduced drug seeking behavior in older HIV-1 Tg animals.

##### Progressive Ratio Responding: Dose Response

Variations in the dose (mg/kg/inf) of cocaine revealed an alteration in the reinforcing efficacy of cocaine in HIV-1 Tg animals treated with SE. Comparison of HIV-1 Tg animals treated with SE and HIV-1 Tg animals treated with vehicle revealed a differential pattern of cocaine self-administration dependent upon dose (Figure 5E)). Specifically, HIV-1 Tg animals treated with vehicle exhibited prominent dose-dependent changes in responding (i.e., a linear (*R*^*2*^= 0.98) decrease in the number of cocaine infusions from the 0.01 to 0.5 mg/kg/inf dose followed by a dramatic increase in the number of cocaine infusions at the 0.75 and 1 mg/kg/inf dose; a dose-dependent effect not observed in HIV-1 Tg animals treated with SE (Best Fit: Horizontal Line). A differential pattern of responding (i.e., HIV-1 Tg SE: Horizontal Line; Control Vehicle: Sigmoidal Dose Response (Variable Slope); *R*^*2*^= 0.99) was also observed when comparing HIV-1 Tg animals treated with SE and control animals treated with vehicle (Figure 5F). Treatment with SE, therefore, precludes the increase in responding at higher cocaine doses supporting a diminished reinforcing efficacy of cocaine in HIV-1 Tg animals. Furthermore, it is notable that utilization of a dose-response experimental paradigm also supports the generalization of reduced drug seeking behavior in older HIV-1 Tg animals across dose.

#### Synaptic Dysfunction in Medium Spiny Neurons of the Nucleus Accumbens

##### Branch Order

Treatment with SE fails to increase dendritic branching complexity in HIV-1 Tg animals in MSNs of the NAc following a history of cocaine self-administration (Figure 6A). Specifically, dendritic branching complexity in HIV-1 Tg animals treated with SE was indistinguishable from HIV-1 Tg animals treated with vehicle (*p*>0.05) and remained significantly decreased relative to control animals treated with vehicle (Genotype x Branch Order interaction with a prominent linear component, [*F*(1,15)=5.3, *p*≤0.04, η^2^p=*0*.259]).

**FIGURE 6.**
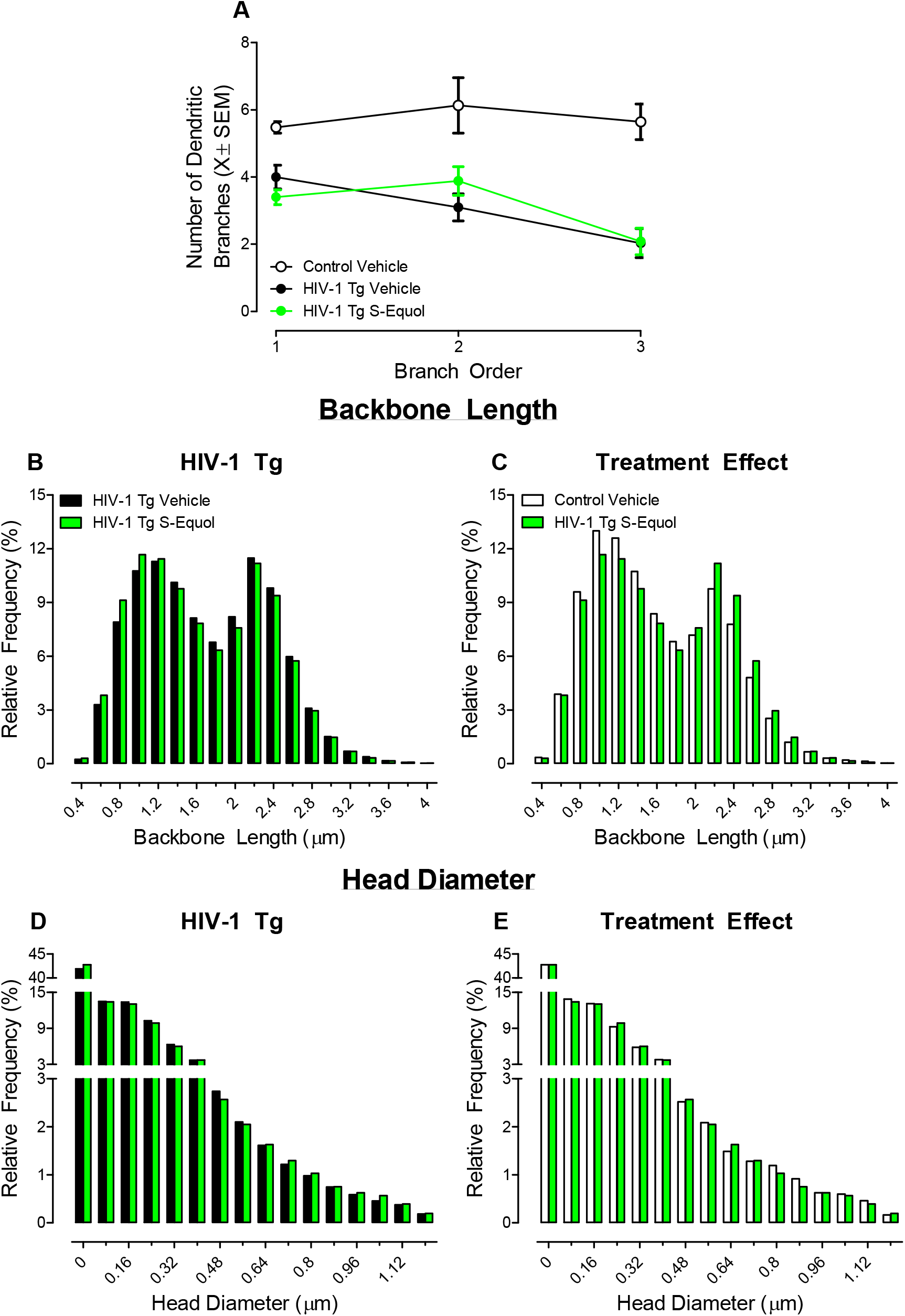
The utility of S-Equol (SE) to modify neuronal (**A**; ±SEM) and dendritic spine morphology (**B-E**) in medium spiny neurons of the nucleus accumbens is illustrated as a function of genotype (HIV-1 Tg vs. Control) and treatment (SE vs. Vehicle; ±95% Confidence Intervals). Although treatment with SE failed to alter dendritic branching (**A**), long-term modification in dendritic spine morphology were observed in HIV-1 Tg animals, evidenced by a prominent population shift towards shorter dendritic spines (**B**) with increased head diameter (**D**) relative to HIV-1 Tg animals treated with vehicle. Treatment with SE shifted the morphological parameters of dendritic spines in MSNs of the NAc to a more mature phenotype relative to HIV-1 Tg animals treated with vehicle, evidenced by a prominent population shift towards shorter dendritic spines (Figure 6B; [*F*(1,1538)=79.6, *p*≤0.01]) with increased head diameter (Figure 6D; [*F*(1,1292)=25.4, *p*≤0.01]). Relative to control animals treated with vehicle, HIV-1 Tg animals treated with SE exhibited significantly longer dendritic spines (**C**), no statistically significant differences in head diameter (**E**) between the two groups were observed.

##### Dendritic Spine Morphology

Morphological parameters, including backbone length and head diameter, of dendritic spines in MSNs of the NAc were enhanced in HIV-1 Tg animals following treated with SE. Treatment with SE shifted the morphological parameters of dendritic spines in MSNs of the NAc to a more mature phenotype relative to HIV-1 Tg animals treated with vehicle, evidenced by a prominent population shift towards shorter dendritic spines (Figure 6B; [*F*(1,1538)=79.6, *p*≤0.01]) with increased head diameter (Figure 6D; [*F*(1,1292)=25.4, *p*≤0.01]). Furthermore, although HIV-1 Tg animals treated with SE exhibited significantly longer dendritic spines (Figure 6C; [*F*(1,1498)=155.5, *p*≤0.01]) relative to control animals treated with vehicle, no statistically significant differences in head diameter between the two groups were observed (Figure 6E; *p*>0.05). Thus, treatment with SE induced a profound population shift in dendritic spine morphology, supporting an enhancement of synaptic efficacy.

## DISCUSSION

Older HIV-1 Tg rats treated with vehicle exhibited prominent alterations in goal-directed behavior relative to their control counterparts; apathetic behavior that was mitigated by treatment with SE. At the genotypic level, apathetic behavior in older HIV-1 Tg rats treated with vehicle was characterized by a diminished reinforcing efficacy of, and sensitivity to, sucrose and enhanced drug seeking for cocaine. Treatment with SE, however, ameliorated alterations in goal-directed behaviors and reduced drug seeking behavior in HIV-1 Tg rats. The therapeutic benefits of SE may be due, at least in part, to the partial restoration of synaptic function, evidenced by a population shift towards a more mature dendritic spine phenotype in HIV-1 Tg animals treated with SE. Taken together, SE restored motivated behavior in older HIV-1 Tg rats, expanding the potential clinical utility of SE to include both neurocognitive and affective alterations.

The assessment of goal-directed behavior, which relies upon Pavlovian conditioning and operant, or instrumental, conditioning [60], was utilized as an index of apathy in the present study.. Pavlovian conditioning utilizes repeated associations of two stimuli to activate behavior. With reference to operant conditioning, the utilization of either “positive” (i.e., the addition of a stimulus following a response) or “negative” (i.e., the removal of a stimulus following a response) reinforcement increases the likelihood that the response will occur again [61]. Once conditioning processes were learned (i.e., via autoshaping acquisition), an aspect of reward expectation (e.g., unit-dose of the reward; schedule of reinforcement: fixed-ratio (FR) vs. progressive ratio (PR)) was manipulated to elucidate changes in goal-directed behavior. Furthermore, the present study examined goal-directed behaviors for both a natural (i.e., sucrose) reinforcer and a drug (i.e., cocaine) reinforcer; reinforcers which uniquely characterize apathetic behaviors in the HIV-1 Tg rat.

Older HIV-1 Tg rats exhibited a diminished reinforcing efficacy of, and sensitivity to, sucrose, as well as an enhanced response vigor for, and greater sensitivity to, cocaine. With regards to sucrose, the behavioral presentation of apathy in the present study is distinct from the decreased response vigor previously reported in young (i.e., 2 months of age) HIV-1 Tg rats [37]. Given the positive correlation between apathy and age in HIV-1 seropositive individuals [16], differences in the behavioral presentation of apathy may be due, at least in part, to age; albeit there were also notable differences in experimental design. With regards to cocaine self-administration, the enhanced response vigor for cocaine in HIV-1 Tg rats on an FR-1 schedule of reinforcement is again distinct from observations by Wayman et al. ([62]; i.e., no statistically significant difference in responding) or Bertrand et al. ([37]; i.e., reduced response vigor); inconsistencies which may reflect differences in age, the experimental protocol, and/or cocaine dose. However, the observed increased sensitivity to cocaine in older HIV-1 Tg rats is consistent with previous observations in relatively young (i.e., 3-4 months of age) HIV-1 Tg rats [63]. Therefore, the differences in sensitivity to cocaine between the Bertrand et al. [37] manuscript and the present study more likely results from differences in cocaine dose. Collectively, independent of differences in the response to either sucrose or cocaine self-administration across studies, the HIV-1 Tg rat exhibits prominent alterations in goal-directed behaviors supporting an advantageous biological system for the evaluation of therapeutics for apathy resulting from chronic HIV-1 viral protein exposure.

Alterations in goal-directed behavior resulting from chronic HIV-1 viral protein exposure, however, were mitigated by treatment with SE; results which extend the therapeutic utility of SE. The therapeutic efficacy of SE, a selective ERβ agonist, for neurocognitive impairments associated with HAND has been critically tested across multiple ages (i.e., treatment beginning at PD 28, 2-3 months of age, and 6-8 months of age), neurocognitive domains (e.g., temporal processing, stimulus-reinforcement learning, sustained attention), and the factor of biological sex [34-36]. Results of the present study extend the therapeutic utility of SE to include the mitigation of apathetic behaviors, as evidenced by the enhanced reinforcing efficacy of, and sensitivity to, sucrose observed in HIV-1 Tg rats treated with SE. Furthermore, treatment with SE dramatically reduced drug seeking behavior in the HIV-1 Tg rat; a finding that deserves further consideration given the well-recognized relationship between estrogens and drug abuse [64] Women are uniquely vulnerable to cocaine addiction, evidenced by a faster acquisition of cocaine self-administration [65-67], greater willingness-to-work for cocaine (i.e., breakpoint; [68]) and a greater sensitivity to cocaine [69-70]; preclinical observations which are consistent with the clinical picture of addiction [71]. The enhanced behavioral responses to cocaine result, at least in part, from the activation of ERβ in the NAc [72]. Notably, when control animals are treated with SE, an agonist selective for ERβ [31], an increased reinforcing efficacy of, and altered sensitivity to, cocaine is observed (Supplementary Figure 1A-C). The structure and function of the NAc, however, is altered by chronic HIV-1 viral protein exposure [e.g., 25-26, 73-74], which may preclude this untoward side effect (i.e., enhanced drug seeking behavior) in HIV-1 Tg rats treated with SE.

Examination of dendritic spine morphology in MSNs of the NAc following a history of cocaine self-administration affords additional evidence for structural alterations in the NAc in the HIV-1 Tg rat. Following a history of cocaine self-administration, HIV-1 Tg animals treated with vehicle exhibited a population shift towards longer dendritic spines with decreased dendritic spine head diameter relative to control animals treated with vehicle; a population shift consistent with a ‘filopodia’-like morphology. However, treatment with SE led to long-term modifications in dendritic spines in MSNs of the NAc in HIV-1 Tg animals. Specifically, HIV-1 Tg animals treated with SE exhibited a prominent population shift towards a more mature phenotype (i.e., ‘thin’) relative to HIV-1 Tg animals treated with vehicle. Given that apathy is mediated in part by the NAc [75-77], the shift in dendritic spine morphology affords a potential mechanism by which SE exerts its therapeutic effects. Critically, the beneficial effects of SE on the morphological parameters of dendritic spines in the HIV-1 Tg rat may reflect the utility of SE to more broadly remodel neuronal circuitry.

Indeed, the ovarian steroid hormone 17β-estradiol acts on multiple brain regions (i.e., PFC, NAc) associated with the anterior cingulate circuit. In the PFC, 17β-estradiol increases dendritic spine density [e.g., 78-81] via the ERβ pathway [81] and induces morphological alterations in dendritic spines [79]. Within the NAc, however, 17β-estradiol significantly reduces dendritic spine density in the NAc core and increases the prevalence of immature spine types (i.e., stubby, filopodia; [82-83]). In a consistent manner, in the present study, control animals treated with SE exhibited a population shift towards a less mature dendritic spine phenotype relative to control animals treated with vehicle (Supplementary Figure 1D-F). To more comprehensively evaluate if and/or how SE remodels neuronal circuitry in the HIV-1 Tg rat, studies in the absence of cocaine and across additional brain regions involved in the anterior cingulate circuit are needed.

In conclusion, following chronic HIV-1 viral protein exposure, prominent alterations in goal-directed behavior and a population shift towards an immature dendritic spine phenotype were observed. In HIV-1 Tg animals, treatment with SE mitigated deficits in goal-directed behavior and had no untoward side effects (e.g., enhancing drug seeking behavior). Furthermore, morphological parameters of dendritic spines in MSNs of the NAc in HIV-1 Tg animals treated with SE were shifted towards a more mature phenotype, supporting a potential mechanism by which SE exerts its therapeutic effects. Taken together, SE restored motivated behavior in the HIV-1 Tg rat, expanding the potential clinical utility of SE to include both neurocognitive and affective alterations.

## ACKNOWLEDGEMENTS

This work was supported in part by grants from NIH (National Institute on Drug Abuse, DA013137; National Institute of Child Health and Human Development HD043680; National Institute of Mental Health, MH106392; National Institute of Neurological Disorders and Stroke, NS100624) and the interdisciplinary research training program supported by the University of South Carolina Behavioral-Biomedical Interface Program.

## AUTHOR CONTRIBUTIONS

R.M.B. and C.F.M. designed the study. S.J.B., C.F.M. and S.B.H. planned the study and S.J.B. and J.M.I. collected the behavioral and neuroanatomical data. K.A.M., S.J.B., J.M.I., and C.F.M. analyzed the data. S.J.B., J.M.I., S.B.H., R.M.B. and C.F.M., wrote the early drafts of the manuscript, and K.A.M., C.F.M., and R.M.B. updated and revised the final draft of the manuscript. All authors critically appraised and approved the final version of the manuscript.

## SUPPLEMENTARY FIGURE 1

**Supplementary Figure S1.**
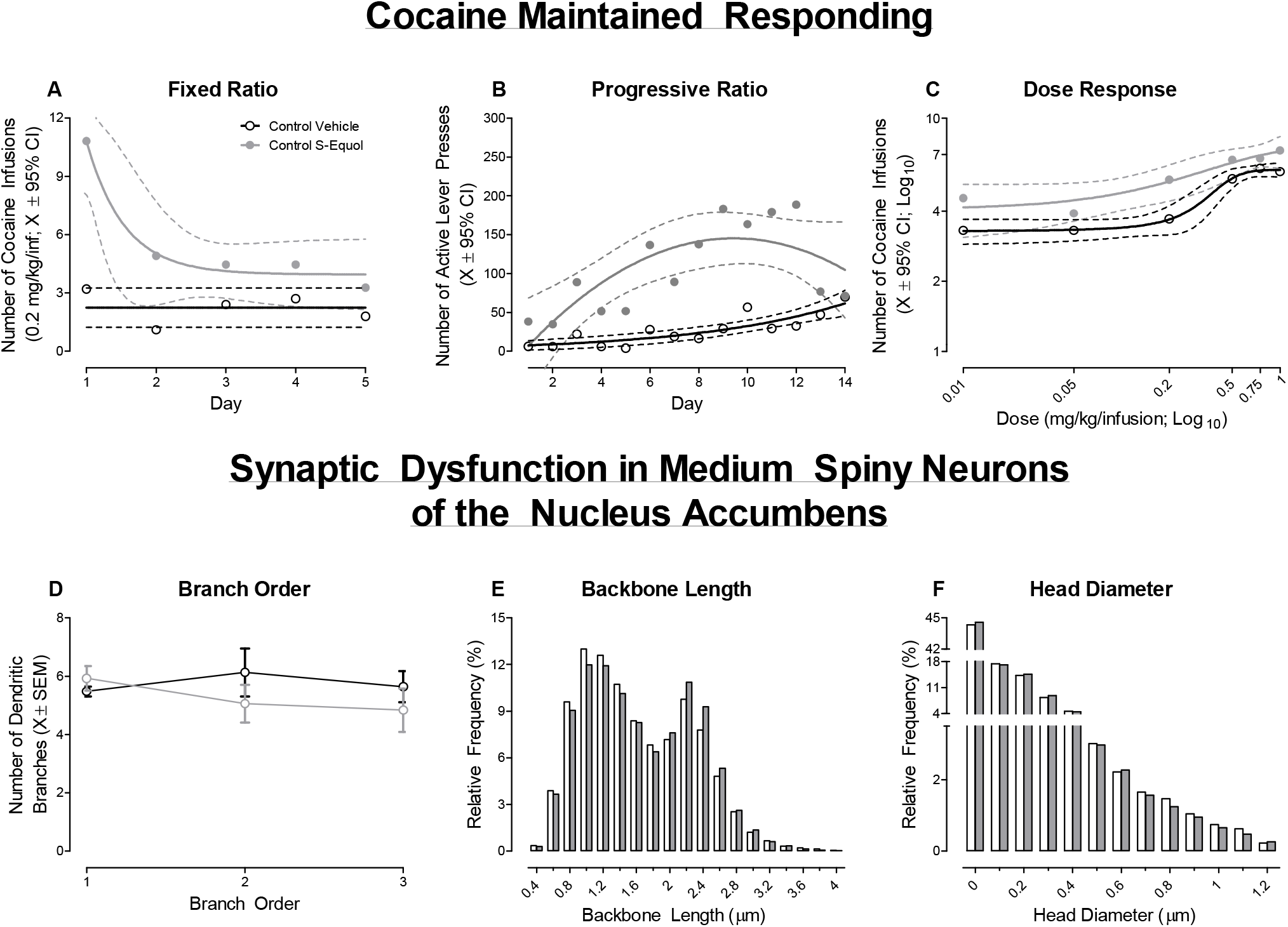
The impact of S-Equol (SE) on cocaine-maintained responding (**A-C**) and medium spiny neurons (MSNs) of the nucleus accumbens (**D-F**) is illustrated for control animals as a function of treatment (Control Vehicle vs. Control SE). Control animals treated with SE exhibited an initial novelty response to cocaine self-administration followed by a rapid decay (**A**). Treatment with SE increased drug seeking behavior in control animals (**B**) and altered the sensitivity to cocaine (**C**). Furthermore, although SE had no significant effect on dendritic branching in MSNs (**D**), profound population shifts towards longer dendritic spines (**E**) with decreased head diameter (**F**) were observed relative to control animals treated with vehicle.

